## Supplementary material for "I knew that! Response-based Outcome Predictions and Confidence Regulate Feedback Processing and Learning": Compiled Supplementary Materials

**Supplement**

**Table S1.** *Follow-up on Prediction and Performance Precision effects on Confidence*

| | Confidence | | | | |
|---|---|---|---|---|---|
| *Predictors* | *Estimates* | *SE* | *CI* | *t* | *p* |
| (Intercept) | 0.26 | 0.04 | 0.18 – 0.33 | 6.36 | **2.055e-10** |
| Block 2-1 | 0.03 | 0.01 | 0.01 – 0.06 | 2.98 | **2.859e-03** |
| Block 3-2 | 0.01 | 0.01 | -0.01 – 0.03 | 0.77 | 4.424e-01 |
| Block 4-3 | 0.03 | 0.01 | 0.01 – 0.06 | 3.03 | **2.424e-03** |
| Block 5-4 | 0.01 | 0.01 | -0.01 – 0.03 | 1.05 | 2.942e-01 |
| Block [1] : SPE | -0.43 | 0.06 | -0.55 – -0.30 | -6.74 | **1.535e-11** |
| Block [2] : SPE | -0.47 | 0.07 | -0.61 – -0.34 | -6.76 | **1.350e-11** |
| Block [3] : SPE | -0.35 | 0.07 | -0.48 – -0.21 | -5.05 | **4.431e-07** |
| Block [4] : SPE | -0.46 | 0.07 | -0.60 – -0.33 | -6.75 | **1.431e-11** |
| Block [5] : SPE | -0.63 | 0.07 | -0.76 – -0.49 | -9.13 | **6.882e-20** |
| Block [1] : Error Magnitude | 0.03 | 0.06 | -0.09 – 0.16 | 0.53 | 5.964e-01 |
| Block [2] : Error Magnitude | 0.15 | 0.07 | 0.01 – 0.29 | 2.14 | **3.219e-02** |
| Block [3] : Error Magnitude | 0.11 | 0.07 | -0.03 – 0.24 | 1.50 | 1.341e-01 |
| Block [4] : Error Magnitude | 0.27 | 0.07 | 0.13 – 0.41 | 3.69 | **2.286e-04** |
| Block [5] : Error Magnitude | 0.26 | 0.07 | 0.11 – 0.40 | 3.51 | **4.561e-04** |
| **Random Effects** | | | **Model Parameters** | | |
| Residuals | 0.12 | | N | 40 | |
| Intercept | 0.06 | | Observations | 9996 | |
| SPE | 0.03 | | log-Likelihood | -3786.661 | |
| Error Magnitude | 0.06 | | Deviance | 7573.321 | |

*Formula: Confidence ~ Block/( Error Magnitude+ SPE) + (Error Magnitude+SPE|participant);*
*Note: ":" indicates interactions*

**Table S2.** *Control analysis for Confidence Calibration effect on Learning*

| | Log Error Magnitude | | | | |
|---|---|---|---|---|---|
| *Predictors* | *Estimates* | *SE* | *CI* | *t* | *p* |
| (Intercept) | 5.05 | 0.06 | 4.93 – 5.18 | 80.64 | **0.000e+00** |
| Confidence Calibration | 0.31 | 0.56 | -0.78 – 1.40 | 0.55 | 5.796e-01 |
| Trial (linear) | -0.46 | 0.07 | -0.59 – -0.33 | -6.83 | **8.486e-12** |
| Trial (quadratic) | 0.12 | 0.02 | 0.07 – 0.16 | 5.03 | **4.784e-07** |
| Response Variance | 0.54 | 0.05 | 0.43 – 0.64 | 9.96 | **2.391e-23** |
| Trial (linear) : Confidence Calibration | -0.66 | 0.31 | -1.27 – -0.05 | -2.12 | **3.393e-02** |
| **Random Effects** | | | **Model Parameters** | | |
| Residuals | 1.17 | | N | 40 | |
| Intercept | 0.11 | | Observations | 9996 | |
| Trial (linear) | 0.03 | | log-Likelihood | -15057.557 | |
| | | | Deviance | 30115.114 | |

*Formula: log Error Magnitude ~ Confidence Calibration*Trial (linear)+ Trial (quadratic)+ Response Variance+ (Trial(linear) |participant); Note: ":" indicates interactions*

**Table S3.** *Follow-up on Block and Confidence effects on Relative Error Signals*

| | Absolute RPE vs EM | | | | RPE vs EM | | | | SPE vs EM | | | |
|---|---|---|---|---|---|---|---|---|---|---|---|---|
| *Predictors* | *Estimates* | *SE* | *t* | *p* | *Estimates* | *SE* | *t* | *p* | *Estimates* | *SE* | *t* | *p* |
| (Intercept) | -40.99 | 3.78 | -10.84 | **2.167e-27** | -115.03 | 8.42 | -13.66 | **1.855e-42** | 1.90 | 3.39 | 0.56 | 5.760e-01 |
| Block | 19.44 | 3.98 | 4.89 | **1.002e-06** | -0.58 | 5.38 | -0.11 | 9.137e-01 | 18.49 | 4.29 | 4.31 | **1.647e-05** |
| Confidence | -25.84 | 9.43 | -2.74 | **6.124e-03** | -7.14 | 19.33 | -0.37 | 7.120e-01 | -54.05 | 9.74 | -5.55 | **2.883e-08** |
| Block : Confidence | -5.29 | 4.94 | -1.07 | 2.844e-01 | 38.44 | 5.28 | 7.28 | **3.334e-13** | -8.24 | 5.61 | -1.47 | 1.418e-01 |
| **Random Effects** | | | | | | | | | | | | |
| Residuals | 16863.59 | | | | 18142.16 | | | | 22090.63 | | | |
| Intercept | 443.78 | | | | 2671.03 | | | | 296.41 | | | |
| Confidence | 2873.45 | | | | 14032.18 | | | | 2952.20 | | | |
| Block | 426.60 | | | | 927.54 | | | | 469.39 | | | |
| N | 40 | | | | 40 | | | | 40 | | | |
| Observations | 9996 | | | | 9996 | | | | 9996 | | | |
| Deviance | 125867.367 | | | | 126731.891 | | | | 128536.108 | | | |
| log-Likelihood | -62933.683 | | | | -63365.945 | | | | -64268.054 | | | |

*Formula: DV ~ Block* Confidence + (Block + Confidence |participant); DVs are Absolute RPE vs Error Magnitude, RPE vs Error Magnitude and SPE vs Error Magnitude; Note: ":" indicates interactions*

**Table S4.** *Follow-up on Confidence by Block Interaction on RPE benefit over Error Magnitude*

| | **RPE vs Error Magnitude** | | | | |
|---|---|---|---|---|---|
| *Predictors* | *Estimates* | *SE* | *CI* | *t* | *p* |
| (Intercept) | -114.67 | 8.41 | -131.16 – -98.18 | -13.63 | **<0.001** |
| Block2-1 | 12.08 | 5.46 | 1.37 – 22.79 | 2.21 | **0.027** |
| Block3-2 | -1.93 | 5.50 | -12.71 – 8.84 | -0.35 | 0.725 |
| Block4-3 | -3.21 | 5.58 | -14.15 – 7.72 | -0.58 | 0.565 |
| Block5-4 | -6.58 | 5.70 | -17.75 – 4.59 | -1.16 | 0.248 |
| Block [1] : Confidence | -46.80 | 20.49 | -86.95 – -6.64 | -2.28 | **0.022** |
| Block [2] : Confidence | -20.33 | 20.25 | -60.01 – 19.35 | -1.00 | 0.315 |
| Block [3] : Confidence | -19.37 | 20.14 | -58.85 – 20.10 | -0.96 | 0.336 |
| Block [4] : Confidence | 15.10 | 20.21 | -24.52 – 54.72 | 0.75 | 0.455 |
| Block [5] : Confidence | 35.03 | 20.46 | -5.07 – 75.14 | 1.71 | 0.087 |
| **Random Effects** | | | | | |
| Residual | 18110.80 | | | | |
| Intercept | 2663.50 | | | | |
| Confidence | 13872.45 | | | | |
| Block | 929.76 | | | | |
| N | 40 | | | | |
| Observations | 9996 | | | | |
| Deviance | 126714.387 | | | | |
| log-Likelihood | -63357.194 | | | | |

*Formula: RPE vs Error Magnitude ~ Block/Confidence +(Confidence|participant);*
*Note: ":" indicates interactions*

**Table S5.** *Block and Confidence effects on Error Signals*

| | Error Magnitude | | | | | RPE | | | | | SPE | | | | |
|---|---|---|---|---|---|---|---|---|---|---|---|---|---|---|---|
| *Predictors* | *Estimates* | *SE* | *CI* | *t* | *p* | *Estimates* | *SE* | *CI* | *t* | *p* | *Estimates* | *SE* | *CI* | *t* | *p* |
| (Intercept) | 204.21 | 10.26 | 184.11 – 224.31 | 19.91 | **3.281e-88** | -91.45 | 9.63 | -110.33 – -72.58 | -9.50 | **2.187e-21** | 206.95 | 10.22 | 186.92 – 226.97 | 20.26 | **3.088e-91** |
| Block | -37.10 | 6.50 | -49.84 – -24.36 | -5.71 | **1.152e-08** | 36.26 | 6.15 | 24.20 – 48.32 | 5.89 | **3.805e-09** | -17.54 | 5.01 | -27.36 – -7.72 | -3.50 | **4.622e-04** |
| Confidence | -16.10 | 13.12 | -41.81 – 9.60 | -1.23 | 2.195e-01 | 18.97 | 10.66 | -1.93 – 39.86 | 1.78 | 7.521e-02 | -71.20 | 8.02 | -86.91 – -55.49 | -8.88 | **6.510e-19** |
| Block : Confidence | 30.09 | 6.74 | 16.87 – 43.30 | 4.46 | **8.110e-06** | -64.48 | 6.92 | -78.06 – -50.91 | -9.31 | **1.240e-20** | 16.99 | 6.08 | 5.07 – 28.90 | 2.79 | **5.198e-03** |
| **Random Effects** | | | | | | | | | | | | | | | |
| Residuals$^2$ | 30497.33 | | | | | 32243.17 | | | | | 26425.43 | | | | |
| Intercept | 3956.08 | | | | | 3442.90 | | | | | 3972.30 | | | | |
| Confidence | 5556.10 | | | | | 3218.71 | | | | | 1572.42 | | | | |
| Block | 1312.50 | | | | | 1113.46 | | | | | 688.02 | | | | |
| N | 40 | | | | | 40 | | | | | 40 | | | | |
| Observations | 9996 | | | | | 9996 | | | | | 9996 | | | | |
| Deviance | 131861.869 | | | | | 132387.369 | | | | | 130376.103 | | | | |
| log-Likelihood | -65930.935 | | | | | -66193.684 | | | | | -65188.052 | | | | |

*Formula: DV ~ Block* Confidence + (Block + Confidence |participant); DVs are Error Magnitude, RPE and SPE; Note: ":" indicates interactions*

**Table S6.** *Follow-up on Block and Confidence effects on Error Signals*

| | Error Magnitude | | | | RPE | | | | SPE | | | |
|---|---|---|---|---|---|---|---|---|---|---|---|---|
| *Predictors* | *Estimates* | *SE* | *t* | *p* | *Estimates* | *SE* | *t* | *p* | *Estimates* | *SE* | *t* | *p* |
| (Intercept) | 203.22 | 10.31 | 19.71 | **1.730e-86** | -90.70 | 9.70 | -9.35 | **8.971e-21** | 206.39 | 10.24 | 20.16 | **2.084e-90** |
| Block2-1 | -67.95 | 6.91 | -9.83 | **8.194e-23** | 54.82 | 7.00 | 7.83 | **4.952e-15** | -46.12 | 6.21 | -7.43 | **1.116e-13** |
| Block3-2 | -4.57 | 6.97 | -0.66 | 5.120e-01 | 6.07 | 7.07 | 0.86 | 3.906e-01 | 4.09 | 6.27 | 0.65 | 5.147e-01 |
| Block4-3 | -3.29 | 7.07 | -0.47 | 6.413e-01 | 5.65 | 7.17 | 0.79 | 4.307e-01 | 4.27 | 6.37 | 0.67 | 5.030e-01 |
| Block5-4 | -10.02 | 7.22 | -1.39 | 1.653e-01 | 16.30 | 7.33 | 2.22 | **2.613e-02** | -8.05 | 6.51 | -1.24 | 2.158e-01 |
| Block [1] * Confidence | -53.92 | 15.96 | -3.38 | **7.263e-04** | 93.07 | 14.41 | 6.46 | **1.047e-10** | -104.54 | 11.53 | -9.07 | **1.231e-19** |
| Block [2] * Confidence | -17.32 | 15.45 | -1.12 | 2.622e-01 | 30.90 | 13.80 | 2.24 | **2.518e-02** | -63.90 | 11.02 | -5.80 | **6.591e-09** |
| Block [3] * Confidence | -9.58 | 15.20 | -0.63 | 5.287e-01 | 24.07 | 13.50 | 1.78 | 7.468e-02 | -57.71 | 10.77 | -5.36 | **8.398e-08** |
| Block [4] * Confidence | 2.66 | 15.34 | 0.17 | 8.621e-01 | -19.50 | 13.66 | -1.43 | 1.534e-01 | -60.16 | 10.93 | -5.50 | **3.734e-08** |
| Block [5] * Confidence | 0.81 | 15.79 | 0.05 | 9.592e-01 | -36.91 | 14.15 | -2.61 | **9.110e-03** | -67.73 | 11.35 | -5.97 | **2.422e-09** |
| **Random Effects** | | | | | | | | | | | | |
| Residuals | 30249.30 | | | | 32101.35 | | | | 26295.02 | | | |
| Intercept | 4001.20 | | | | 3497.82 | | | | 3989.42 | | | |
| Confidence | 5509.85 | | | | 3396.66 | | | | 1545.22 | | | |
| Block | 1260.21 | | | | 1077.63 | | | | 650.86 | | | |
| N | 40 | | | | 40 | | | | 40 | | | |
| Observations | 9996 | | | | 9996 | | | | 9996 | | | |
| Deviance | 131780.444 | | | | 132344.613 | | | | 130325.232 | | | |
| log-Likelihood | -65890.222 | | | | -66172.306 | | | | -65162.616 | | | |

*Formula: DV ~ Block/Confidence + (Block + Confidence |participant); DVs are Error Magnitude, RPE and SPE; Note: ":" indicates interactions*

**Table S7.** *Block and Confidence effects on Error Signals*

| | **RPE** | | | | **SPE** | | | |
|---|---|---|---|---|---|---|---|---|
| *Predictors* | *Estimates* | *SE* | *t* | *p* | *Estimates* | *SE* | *t* | *p* |
| (Intercept) | -82.91 | 7.46 | -11.11 | **1.085e-28** | 201.62 | 5.16 | 39.04 | **0.000e+00** |
| Block | 10.30 | 4.81 | 2.14 | **3.228e-02** | 3.65 | 3.34 | 1.09 | 2.736e-01 |
| Confidence | 9.13 | 15.52 | 0.59 | 5.561e-01 | -60.12 | 6.68 | -9.00 | **2.255e-19** |
| Error Magnitude | -728.76 | 7.24 | -100.67 | **0.000e+00** | 572.09 | 7.27 | 78.71 | **0.000e+00** |
| Block : Confidence | -45.59 | 4.94 | -9.23 | **2.641e-20** | 0.10 | 4.76 | 0.02 | 9.839e-01 |
| **Random Effects** | | | | | | | | |
| Residuals | 15947.32 | | | | 16472.17 | | | |
| Intercept | 2080.51 | | | | 934.32 | | | |
| Confidence | 8858.40 | | | | 1160.03 | | | |
| Block | 720.24 | | | | 243.98 | | | |
| N | 40 | | | | 40 | | | |
| Observations | 9996 | | | | 9996 | | | |
| Deviance | 125421.104 | | | | 125603.380 | | | |
| log-Likelihood | -62710.552 | | | | -62801.690 | | | |

*Formula: DV ~ Block* Confidence + Error Magnitude, + (Block + Confidence |participant);*
*DVs are RPE and SPE; Note: ":" indicates interactions*

**Table S8.** *Follow-up analyses on Confidence-weighted Predicted Error Magnitude effects on P3b*

| | P3b Amplitude | | | | |
|---|---|---|---|---|---|
| *Predictors* | *Estimates* | *SE* | *CI* | *t* | *p* |
| (Intercept) | 4.27 | 0.30 | 3.68 – 4.85 | 14.28 | **2.949e-46** |
| Block2-1 | -0.23 | 0.23 | -0.69 – 0.22 | -1.01 | 3.146e-01 |
| Block3-2 | -0.10 | 0.23 | -0.56 – 0.35 | -0.43 | 6.652e-01 |
| Block4-3 | -0.31 | 0.24 | -0.77 – 0.15 | -1.31 | 1.886e-01 |
| Block5-4 | 0.08 | 0.24 | -0.39 – 0.54 | 0.32 | 7.472e-01 |
| Error Magnitude | -1.04 | 0.50 | -2.01 – -0.07 | -2.09 | **3.652e-02** |
| Sensory Prediction Error | 1.50 | 0.40 | 0.71 – 2.29 | 3.73 | **1.944e-04** |
| Block [1] : Confidence | 1.23 | 0.38 | 0.48 – 1.98 | 3.23 | **1.225e-03** |
| Block [2] : Confidence | 0.40 | 0.36 | -0.30 – 1.10 | 1.13 | 2.569e-01 |
| Block [3] : Confidence | 0.02 | 0.36 | -0.67 – 0.72 | 0.07 | 9.473e-01 |
| Block [4] : Confidence | -0.09 | 0.35 | -0.78 – 0.60 | -0.26 | 7.945e-01 |
| Block [5] : Confidence | 0.02 | 0.35 | -0.67 – 0.70 | 0.05 | 9.634e-01 |
| Block [1] : Predicted Error Magnitude | -0.14 | 0.96 | -2.03 – 1.75 | -0.14 | 8.874e-01 |
| Block [2] : Predicted Error Magnitude | 0.52 | 1.00 | -1.43 – 2.47 | 0.52 | 5.999e-01 |
| Block [3] : Predicted Error Magnitude | -0.74 | 1.06 | -2.83 – 1.35 | -0.69 | 4.878e-01 |
| Block [4] : Predicted Error Magnitude | -1.70 | 0.97 | -3.60 – 0.20 | -1.76 | 7.901e-02 |
| Block [5] : Predicted Error Magnitude | -2.13 | 0.82 | -3.74 – -0.52 | -2.59 | **9.493e-03** |
| Block [1] : Confidence : Predicted Error Magnitude | -5.39 | 1.52 | -8.37 – -2.41 | -3.55 | **3.859e-04** |
| Block [2] : Confidence : Predicted Error Magnitude | -1.63 | 1.54 | -4.66 – 1.39 | -1.06 | 2.895e-01 |
| Block [3] : Confidence : Predicted Error Magnitude | -1.60 | 1.67 | -4.86 – 1.67 | -0.96 | 3.381e-01 |
| Block [4] : Confidence : Predicted Error Magnitude | 3.17 | 1.44 | 0.34 – 6.00 | 2.20 | **2.796e-02** |
| Block [5] : Confidence : Predicted Error Magnitude | 0.83 | 1.16 | -1.43 – 3.10 | 0.72 | 4.721e-01 |
| **Random Effects** | | | | | |
| Residuals | 23.93 | | | | |
| Intercept | 3.27 | | | | |
| Error Magnitude | 3.45 | | | | |
| Confidence | 0.74 | | | | |
| N | 40 | | | | |
| Observations | 9678 | | | | |
| Deviance | 58386.274 | | | | |
| log-Likelihood | -29193.137 | | | | |

*Formula: P3b ~ Block/(Confidence*Predicted Error Magnitude+ SPE)+Error Magnitude + (Error Magnitude+Confidence|participant); Note: ":" indicates interactions*

**Table S9.** *Trial to Trial Improvements by Block and Previous Error and modulations by Previous P3b*

| | **Improvement** | | | | | **Improvement** | | | | |
|---|---|---|---|---|---|---|---|---|---|---|
| *Predictors* | *Estimates* | *SE* | *CI* | *t* | *p* | *Estimates* | *SE* | *CI* | *t* | *p* |
| (Intercept) | -164.33 | 9.53 | -183.00 – -145.66 | -17.25 | **1.102e-66** | -163.29 | 9.78 | -182.46 – -144.12 | -16.69 | **1.491e-62** |
| Block | -0.90 | 3.75 | -8.25 – 6.44 | -0.24 | 8.094e-01 | -3.27 | 4.88 | -12.83 – 6.28 | -0.67 | 5.019e-01 |
| Error Magnitude (n-1) | 0.85 | 0.01 | 0.83 – 0.87 | 84.55 | **0.000e+00** | 0.84 | 0.01 | 0.81 – 0.86 | 68.75 | **0.000e+00** |
| Block : Error Magnitude (n-1) | 0.13 | 0.01 | 0.11 – 0.15 | 10.47 | **1.159e-25** | 0.15 | 0.02 | 0.12 – 0.18 | 9.84 | **7.849e-23** |
| P3b (n-1) | | | | | | -0.87 | 3.51 | -7.75 – 6.00 | -0.25 | 8.036e-01 |
| P3b (n-1) : Error Magnitude (n-1) | | | | | | 0.01 | 0.01 | -0.01 – 0.03 | 0.88 | 3.765e-01 |
| Block : P3b (n-1) | | | | | | 4.34 | 4.86 | -5.18 – 13.86 | 0.89 | 3.715e-01 |
| Block : P3b (n-1) : Error Magnitude (n-1) | | | | | | -0.03 | 0.01 | -0.06 – -0.00 | -2.16 | **3.058e-02** |
| **Random Effects** | | | | | | | | | | |
| Residual | 37789.84 | | | | | 36777.96 | | | | |
| Intercept | 3327.45 | | | | | 3324.68 | | | | |
| N | 40 | | | | | 40 | | | | |
| Observations | 9956 | | | | | 9638 | | | | |
| Deviance | 133313.382 | | | | | 128797.375 | | | | |
| log-Likelihood | -66656.691 | | | | | -64398.688 | | | | |

*Formula: Improvement ~ Block* Previous Error Magnitude + (1|participant); Improvement ~ Block* Previous Error Magnitude*Previous P3b + (1|participant)*
*Note: ":" indicates interactions*

**Table S10.** *Follow-up on Trial to Trial Improvements by Block and Previous Error and modulations by Previous P3b*

| | Improvement | | | | | Improvement | | | | |
|---|---|---|---|---|---|---|---|---|---|---|
| *Predictors* | *Estimates* | *SE* | *CI* | *t* | *p* | *Estimates* | *SE* | *CI* | *t* | *p* |
| (Intercept) | -167.22 | 9.69 | -186.21 – -148.23 | -17.26 | **9.611e-67** | -166.61 | 9.91 | -186.04 – -147.18 | -16.81 | **2.155e-63** |
| fblock2-1 | -17.90 | 8.37 | -34.30 – -1.49 | -2.14 | **3.251e-02** | -27.65 | 11.14 | -49.48 – -5.83 | -2.48 | **1.303e-02** |
| fblock3-2 | 10.61 | 8.54 | -6.13 – 27.34 | 1.24 | 2.141e-01 | 22.74 | 11.17 | 0.84 – 44.64 | 2.04 | **4.183e-02** |
| fblock4-3 | 1.24 | 8.53 | -15.48 – 17.97 | 0.15 | 8.841e-01 | -8.19 | 11.00 | -29.75 – 13.38 | -0.74 | 4.568e-01 |
| fblock5-4 | 11.99 | 8.43 | -4.54 – 28.52 | 1.42 | 1.552e-01 | 15.11 | 10.68 | -5.83 – 36.05 | 1.41 | 1.572e-01 |
| Block[1] * EM (n-1) | 0.66 | 0.02 | 0.63 – 0.69 | 43.37 | **0.000e+00** | 0.62 | 0.02 | 0.59 – 0.66 | 33.62 | **7.714e-248** |
| Block[2] * EM (n-1) | 0.94 | 0.02 | 0.90 – 0.99 | 43.44 | **0.000e+00** | 0.96 | 0.03 | 0.91 – 1.01 | 36.48 | **2.119e-291** |
| Block[3] * EM (n-1) | 0.92 | 0.02 | 0.87 – 0.96 | 38.53 | **0.000e+00** | 0.89 | 0.03 | 0.84 – 0.95 | 29.77 | **9.296e-195** |
| Block[4] * EM (n-1) | 0.91 | 0.02 | 0.86 – 0.95 | 40.68 | **0.000e+00** | 0.91 | 0.03 | 0.86 – 0.97 | 32.33 | **2.862e-229** |
| Block[5] * EM (n-1) | 0.90 | 0.02 | 0.85 – 0.95 | 37.24 | **1.711e-303** | 0.90 | 0.03 | 0.85 – 0.96 | 31.28 | **7.981e-215** |
| Block[1] * P3b (n-1) | | | | | | -8.39 | 7.85 | -23.78 – 7.00 | -1.07 | 2.854e-01 |
| Block[2] * P3b (n-1) | | | | | | 11.45 | 7.72 | -3.69 – 26.58 | 1.48 | 1.382e-01 |
| Block[3] * P3b (n-1) | | | | | | -8.10 | 7.92 | -23.62 – 7.42 | -1.02 | 3.063e-01 |
| Block[4] * P3b (n-1) | | | | | | 4.19 | 7.71 | -10.92 – 19.31 | 0.54 | 5.865e-01 |
| Block[5] * P3b (n-1) | | | | | | 0.81 | 7.59 | -14.07 – 15.69 | 0.11 | 9.151e-01 |
| Block[1] * P3b (n-1) * EM (n-1) | | | | | | 0.06 | 0.02 | 0.03 – 0.10 | 3.35 | **8.209e-04** |
| Block[2] * P3b (n-1) * EM (n-1) | | | | | | -0.04 | 0.03 | -0.09 – 0.01 | -1.48 | 1.389e-01 |
| Block[3] * P3b (n-1) * EM (n-1) | | | | | | 0.02 | 0.03 | -0.04 – 0.08 | 0.71 | 4.805e-01 |
| Block[4] * P3b (n-1) * EM (n-1) | | | | | | -0.01 | 0.03 | -0.06 – 0.04 | -0.38 | 7.030e-01 |
| Block[5] * P3b (n-1) * EM (n-1) | | | | | | -0.01 | 0.03 | -0.07 – 0.05 | -0.31 | 7.566e-01 |

| | | |
|---|---|---|
| **Random Effects** | | |
| Residual | 37335.51 | 36296.90 |
| Intercept | 3453.25 | 3427.28 |
| N | 40 | 40 |
| Observations | 9956 | 9638 |
| Deviance | 133194.844 | 128672.144 |
| log-Likelihood | -66597.422 | -64336.072 |

*Formula: Improvement ~ Block/Previous Error Magnitude + (1|participant); Improvement ~ Block/(Previous Error Magnitude*Previous P3b) + (1|participant)*
*Note: ":" indicates interactions*

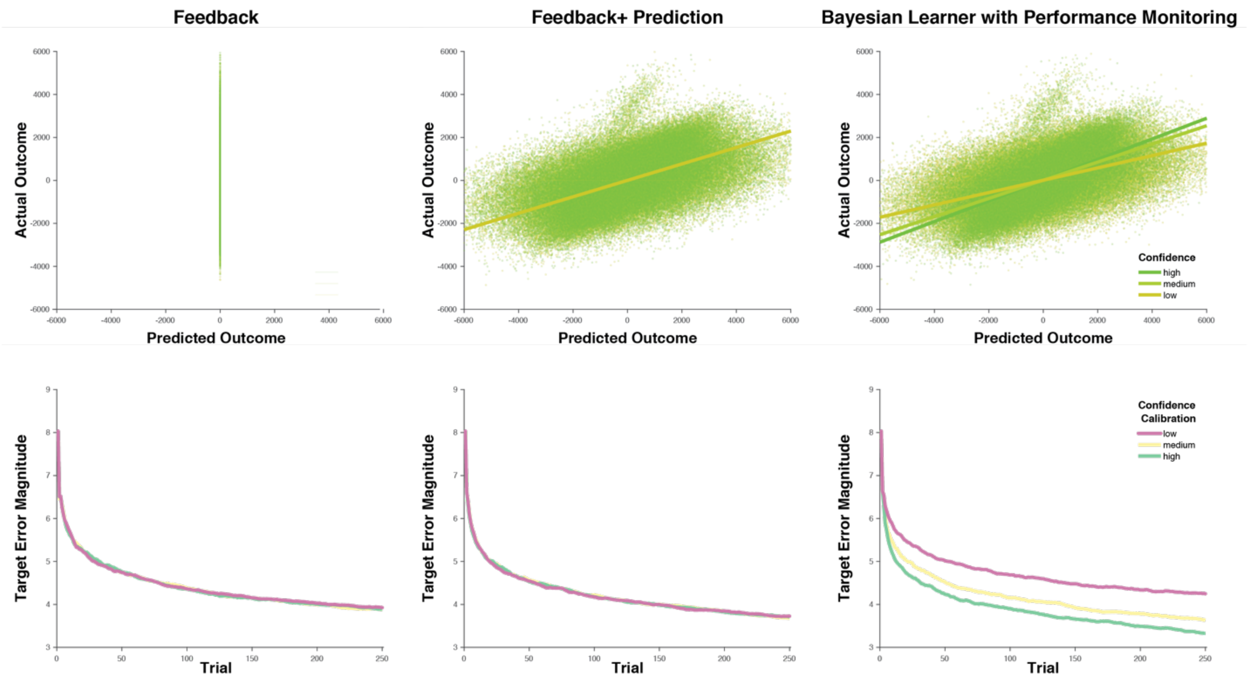


*Figure S 1.* **Model Comparison**. Shown are the relationship between predicted and actual outcomes as a function of Confidence (top) and the Target Error Magnitude as a function of Confidence Calibration for the model that learns from feedback only (left), the model that learns from feedback and outcome predictions (center) and our Bayesian Learner with Performance Monitoring. While the Feedback only model learns the target adequately, it is not able to predict outcomes of its actions. The Feedback and Prediction model learns faster and is able to predict outcomes, however, it cannot distinguish between accurate and inaccurate predictions. Finally, the Bayesian Learner with performance monitoring predicts outcomes and distinguishes between accurate and inaccurate predictions. Whether these are advantageous depends on the fidelity of confidence as a read-out of the precision of the predictions, i.e. confidence calibration. The well calibrated Bayesian Learner with Performance Monitoring outperforms both alternative models.

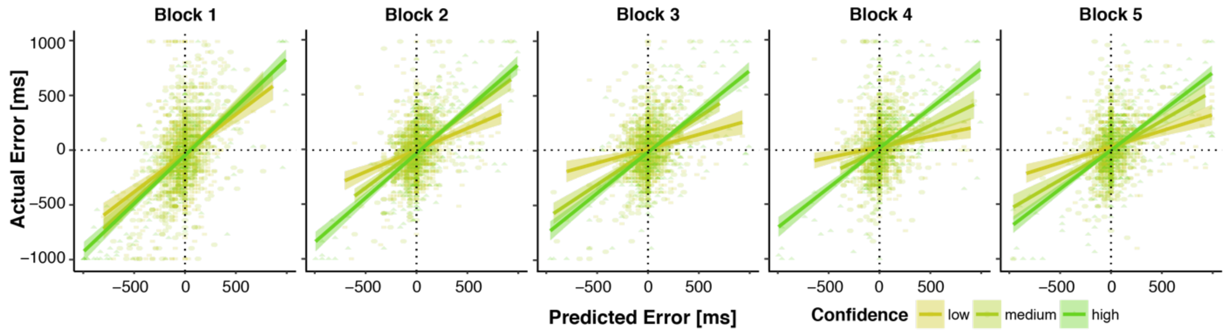


*Figure S 2.***Predictions and Confidence improve as learning progresses.** Plotted are actual errors as a function of predicted errors and confidence terciles. Regression lines represent local linear models. In block 1, many large actual errors are inaccurately predicted to be zero or small. These prediction errors decrease over time. Across blocks, Confidence further dissociates increasingly well between accurate and inaccurate predictions.

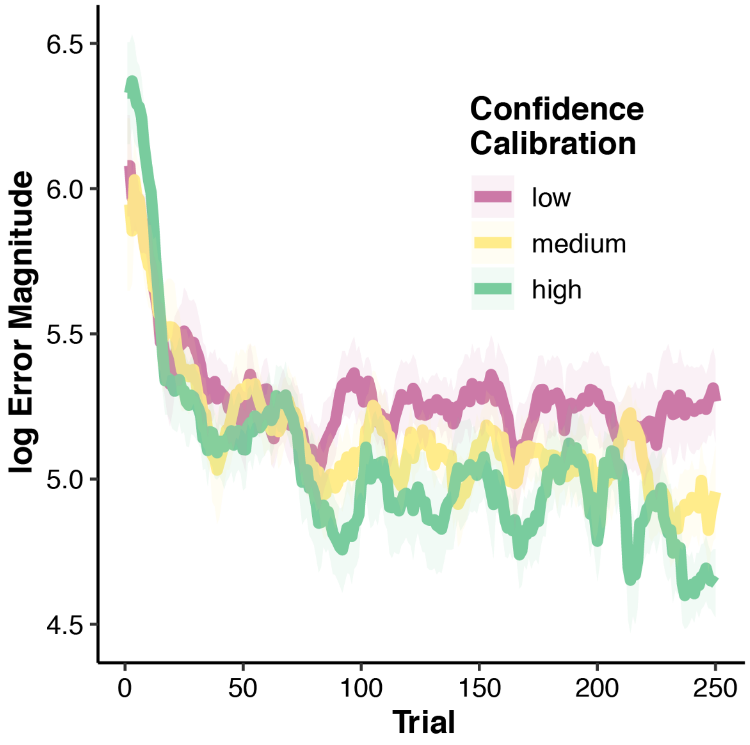


*Figure S 3.* **Running average log error magnitude across trials.** Running average performance averaged across participants within terciles of Confidence calibration. Shaded error bars represent standard error of the mean.

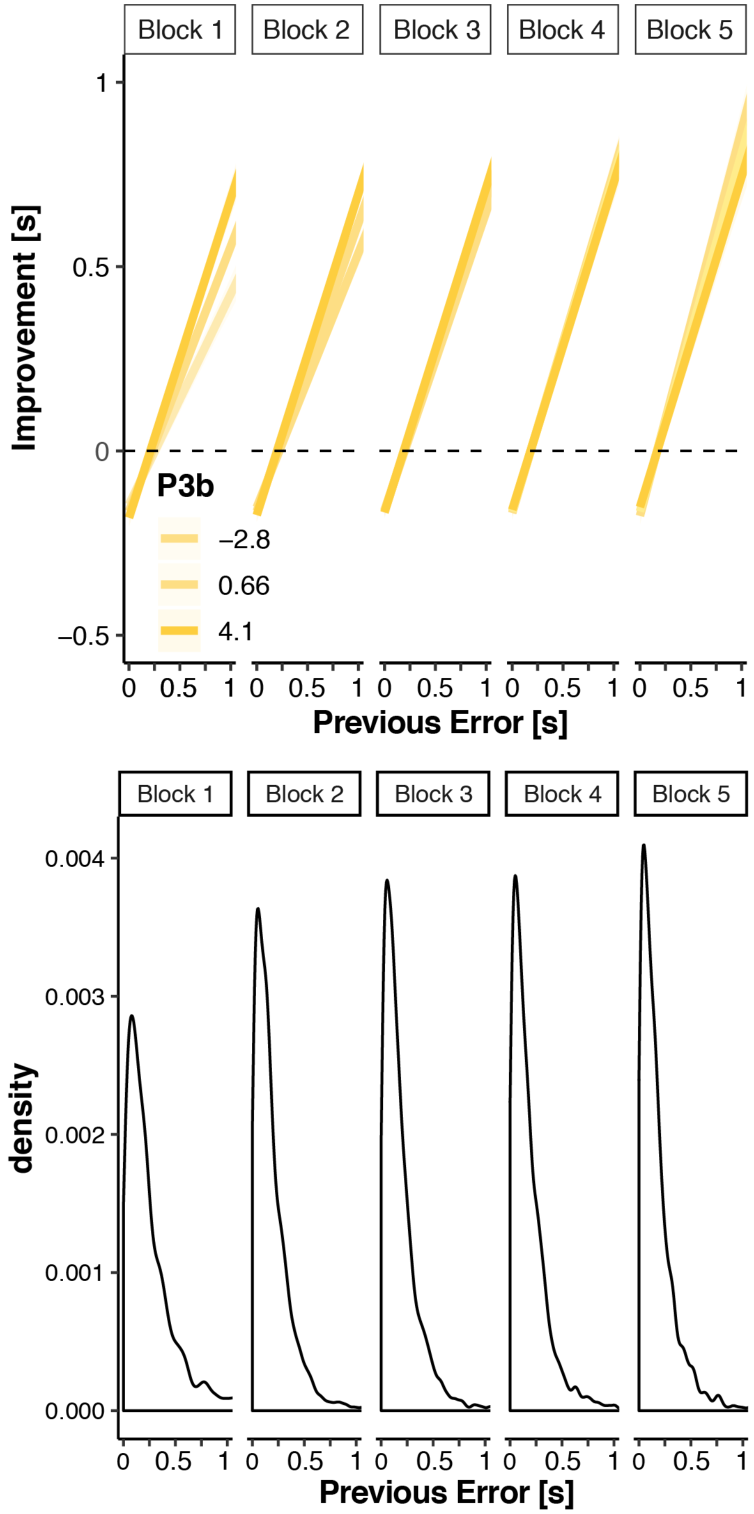


*Figure S 4.* **P3b to feedback modulates error-related adjustments on the subsequent trial.** Top: y-hats illustrating the interaction of previous error magnitude and P3b on improvement on the current trial for each block. Bottom: Distribution of errors in each block.
